# VIP interneurons control hippocampal place cell remapping through transient disinhibition in novel environments

**DOI:** 10.1101/2025.02.01.636072

**Authors:** Máté Neubrandt, Nora Lenkey, Koen Vervaeke

## Abstract

Hippocampal place cells dynamically reorganize their activity, or “remap”, to encode novel spatial environments, a process tightly regulated by inhibitory networks. Here, we identify vasoactive intestinal peptide (VIP)-expressing interneurons as key regulators of this plasticity. Using all-optical methods in mice navigating virtual reality, we show that novel environments trigger a transient increase in VIP interneuron activity, which facilitates rapid place field formation and stable spatial map construction. Bidirectional optogenetic manipulation —suppressing or upregulating VIP interneuron activity— modulates the speed of spatial map formation and alters signatures of behavioral timescale plasticity (BTSP), a mechanism critical for place field induction. Both insufficient and excessive VIP interneuron activity impair reward-seeking behavior specifically in novel environments, revealing that a precise balance of VIP activity is essential for spatial learning. Our findings demonstrate that VIP interneurons gate a temporal window for place cell plasticity, mediating the impact of environmental novelty on hippocampal remapping and spatial memory.

## Introduction

The hippocampus plays a central role in the formation of spatial memories and navigation, with place cells forming dynamic cognitive maps of an animal’s environment ^1,2^. These maps are not static; they undergo rapid reorganization, termed “remapping”, when animals encounter novel surroundings ^3–7^, enabling the brain to adapt to new experiences ^8,9^. Remarkably, this remapping can occur within seconds ^7,10–14,^ underscoring the brain’s extraordinary plasticity. While the environmental cues that trigger remapping have been extensively studied, the specific cellular and circuit mechanisms that enable the rapid formation of new place fields in novel environments remain elusive. In particular, the role of inhibitory interneurons in gating this process is not well understood.

Neuronal inhibition critically shapes place cell activity, regulating not only whether a place cell fires but also the precise timing and rate of firing ^15–18^. While balanced excitation and inhibition in cortical circuits typically maintain stable representations ^19–21^, the rapid formation of new spatial maps in novel environments suggests a more dynamic role for inhibition. Early hippocampal recordings hinted at this, revealing a transient decrease in inhibitory neuron firing while exploring new spaces ^4,5,22,23^. Subsequent studies pinpointed specific interneuron subtypes in hippocampal area CA1 — those expressing parvalbumin (PV) and somatostatin (SOM) — that exhibit this reduced activity during novelty ^24–26^. Moreover, artificially silencing inhibitory neurons promotes the formation of new place fields ^27,28^. Together, these findings suggest the existence of an ‘inhibitory gating’ mechanism that facilitates rapid remapping of place cells in novel environments. However, the specific interneuron subtypes and circuit mechanisms underlying this process remain unknown.

We hypothesized that vasoactive intestinal peptide (VIP)-expressing interneurons, a unique class of inhibitory cells known for their disinhibitory effects ^29– 34^, play a critical role in gating place cell remapping in novel environments. VIP neurons are ideally positioned to regulate dendritic integration in pyramidal cells: They preferentially target SOM interneurons ^26,30,35^, which suppress the distal dendrites of CA1 pyramidal neurons ^17,36–39^. These distal dendrites are critical for dendritic spikes that drive behavioral timescale plasticity (BTSP), a mechanism critical for place field formation ^11,12,40,41^.

Several lines of evidence support our hypothesis that VIP interneurons regulate remapping. Recent work by Tamboli et al. ^42^ demonstrated that hippocampal VIP neurons detect environmental novelty, but their role in place cell remapping and spatial learning was not determined. Furthermore, VIP neurons are engaged by locomotion ^26,35,43–50^, active sensing ^51–53^, and attention ^54,55^, behaviors tightly linked to spatial exploration and learning. They are also potently excited by acetylcholine ^56,57^, a neuromodulator released during arousal and exploration. Here, we tested the hypothesis that VIP interneurons gate place cell remapping in novel environments. Using virtual reality and all-optical methods in mice, we found that VIP neuron activity transiently increases during novel environment exploration. Optogenetic silencing of VIP interneurons delayed the formation of stable place maps, while enhancing their activity accelerated it. However, both manipulations impaired features of BTSP and spatial learning, revealing that a precise balance of VIP interneuron activity is essential for efficient encoding of new environments.

## Results

### Environmental novelty drives parallel increases in CA1 VIP neuronal activity and arousal

To investigate the role of VIP neurons in encoding novel environments, we trained VIP-Cre mice ^58^ to navigate a virtual reality (VR) environment while head-fixed on a treadmill (**Figure 1A**). After 1-2 weeks of training in a familiar environment, mice were exposed to a novel environment in a single session consisting of 30 trials in the familiar environment, 60 trials in the novel environment, and 20 trials back in the familiar environment (**Figure 1A, B**). In the familiar environment, mice exhibited anticipatory behaviors, such as slowing down and increased licking near the hidden reward zones, indicating successful learning of reward locations. In the novel environment, these anticipatory behaviors were significantly reduced, suggesting impaired reward location prediction (**Figure 1C**, 6 mice, 6 sessions).

**Figure 1.**
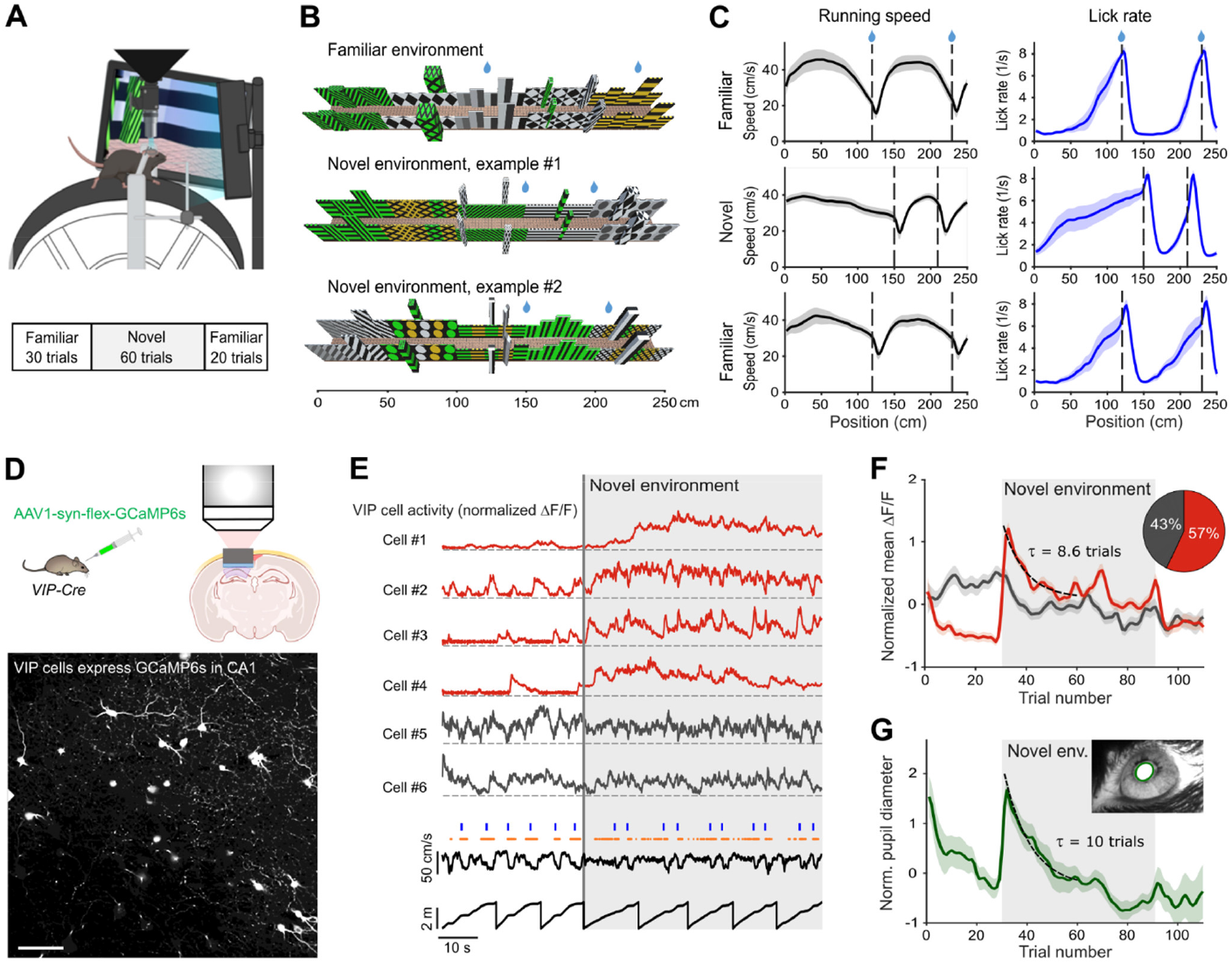
Environmental novelty drives parallel increases in CA1 VIP neuronal activity and arousal. **A)** Top: Cartoon of the experimental setup. A head-fixed mouse runs on a treadmill while navigating a virtual reality (VR) environment under a two-photon microscope. Bottom: Trial block structure of a session, showing transitions between familiar and novel environments. **B)** Familiar VR environment (reward locations: 120 and 230 cm) and two example novel environments (reward locations: 150 and 200 cm or 150 and 210 cm). **C)** Mean running speed (black) and lick rate (blue) during a transition from the familiar to the novel environment, and back to the familiar environment (6 mice, 6 sessions, mean ± SEM across mice). **D)** Example two-photon image of VIP neurons expressing GCaMP6s in the pyramidal layer of hippocampal CA1. Scale bar: 100 µm. **E)** Top: Fluorescence traces (DF/F) of 6 VIP neurons during the transition from familiar to novel environments (gray line). Red: VIP neurons with significantly increased activity; black: unmodulated VIP neurons. Significance was determined by comparing the mean DF/F of the last 10 familiar trials to the first 10 novel trials (paired t-test, one-sided, p < 0.05). Bottom: Running speed and track position traces. Blue ticks: reward delivery; red ticks: licks. **F)** Average VIP neuronal activity (z-scored DF/F) during transitions between familiar and novel environments (6 mice, 149 VIP cells from 10 sessions; mean ± SEM). Red: 85 upregulated VIP cells; black: 64 unmodulated VIP cells. Pie chart shows the proportion of each category. Dashed line: single exponential fit to the decay phase of positively modulated VIP cells. **G)** Simultaneously recorded changes in pupil diameter (z-scored, 5 mice, 8 sessions; mean ± SEM). Dashed line: single exponential fit.

To measure VIP interneuron activity in hippocampal area CA1, we expressed the calcium indicator GCaMP6s ^59^ in VIP-Cre mice and used two-photon microscopy to image their activity (**Figure 1D**). Upon transition to the novel environment, most VIP neurons increased their activity, with some showing rapid onset and others showing a more gradual increase over several trials (**Figure 1E**). Overall, 57% of VIP interneurons increased their activity in the novel environment, while 43% were unmodulated (**Figure 1F**; 149 VIP cells imaged in total, 6 mice, 10 sessions; one field of view (FOV) imaged per session). The mean response of positively modulated neurons showed a rapid increase followed by a slow decrease over the ensuing tens of trials (**Figure 1F**; decay time constant, τ = 8.6 trials), returning to baseline as mice re-entered the familiar environment.

Because VIP neurons are particularly responsive to neuromodulators released during enhanced arousal ^56,60^, we also measured pupil diameter changes as a proxy for arousal when mice entered the novel environment. Pupil diameter mirrored the mean VIP interneuron response, showing a similar rapid increase followed by a slow decrease (**Figure 1G;** decay time constant, τ = 10 trials; 5 mice, 8 sessions). This correlation was also evident at the session onset, where increased pupil diameter coincided with elevated VIP activity (**Figure 1F, G)**. These data demonstrate that VIP interneuron activity increases during novelty exposure, with dynamics paralleling pupil diameter changes, reflecting heightened arousal.

### VIP neurons control the formation of place cells in novel environments

To investigate how VIP interneurons influence place cell remapping, we combined *in vivo* two-photon Ca^2+^ imaging with optogenetic manipulations using VIP-Cre x Thy1-GCaMP6s mice ^58,61^. This allowed us to simultaneously monitor the activity of Thy1-positive excitatory neurons (place cells) while selectively manipulating VIP interneuron activity using the red-shifted opsin ArchT ^62^ for inhibition or ChrimsonR ^63^ for excitation (**Figure 2A, B**). We refer to our previous work for a thorough validation of this approach ^64^, which includes *in vitro* patch-clamp characterization of VIP neurons expressing ArchT and ChrimsonR, *in vivo* characterization using simultaneous expression of GcaMP6s and Chrimson in VIP cells, and histological analysis of VIP cells expressing these opsins.

**Figure 2.**
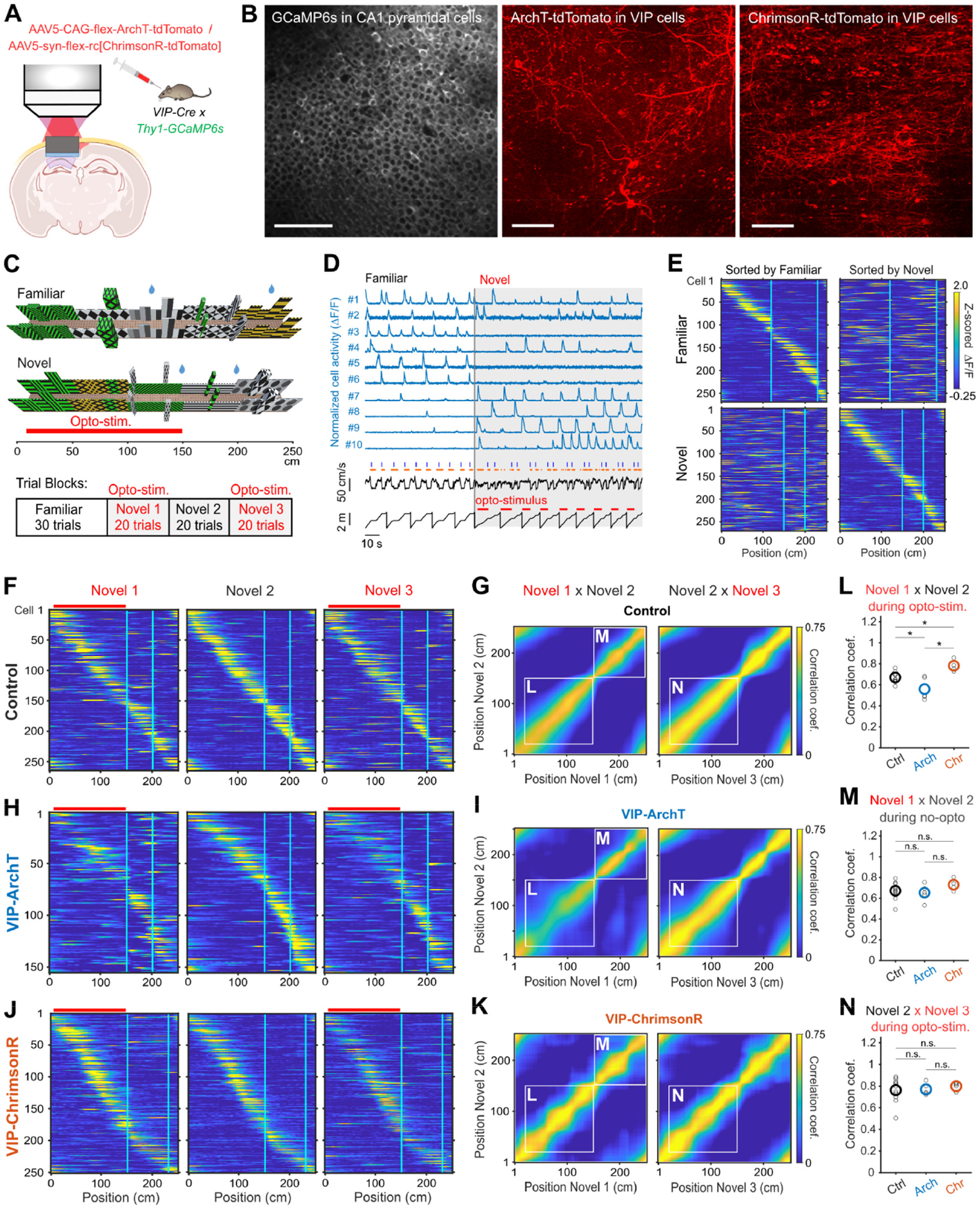
VIP neurons control the formation of place cells in novel environments. **A)** Experimental design: Simultaneous two-photon imaging of CA1 pyramidal neurons expressing GCaMP6s and optogenetic manipulation of VIP neurons expressing the red-shifted inhibitory opsin ArchT or the excitatory opsin ChrimsonR. **B)** Example in vivo two-photon microscope images showing example fields of view of GCaMP6s expression in CA1 pyramidal cells (left), ArchT expression in VIP neurons (middle), and ChrimsonR expression in VIP neurons (right) in VIP neurons. Scale bars: 100 µm. **C)** Familiar and novel VR-environments. Optogenetic stimulation was restricted to the first 1.5 m of the track, up to the first reward location. Bottom: trial block structure within recording session. **D)** Example activity (DF/F) traces of 10 place cells during the transition from familiar to novel environments (gray line). Bottom: running speed and track position traces. Blue ticks: reward delivery; red ticks: licks. **E)** Example session showing place cell remapping in control mice. Top: Place cells in the familiar environment, sorted by peak activity. Bottom: Place cells in the novel environment, sorted by peak activity. Vertical lines indicate the reward locations. **(F, H** and **J)** Example sessions showing the activity of the same place cells across ‘Novel 1’, ‘Novel 2’, and ‘Novel 3’ trial blocks. Top: Control mice. Middle: VIP-ArchT mice. Bottom: VIP-ChrimsonR mice. Place cells sorted by peak activity in the non-stimulated ‘Novel 2’ block. Same color scale as in (E). Vertical lines indicate the reward locations. Red bars indicate optogenetic stimulus location. **(G, I** and **K)** Mean population vector correlations of pyramidal cell activity between ‘Novel 1’ vs. ‘Novel 2’ (left), and ‘Novel 2’ vs. ‘Novel 3’ (right). Control: 6 mice, 11 sessions, VIP-ArchT: 5 mice, 5 sessions, VIP-ChrimsonR: 4 mice, 4 sessions. White boxes: Track segments used for correlation analysis in (L–N). **L)** Population vector correlations using optogenetically stimulated track segments (‘Novel 1’ vs. ‘Novel 2’, based on data in G, I, K). Unpaired two-tailed *t*-tests; Control vs. VIP-ArchT (p = 0.012), Control vs. VIP-ChrimsonR (p = 0.0028) and VIP-ArchT vs. VIP-ChrimsonR (p = 0.0072). **M)** Population vector correlations using track segments without optogenetic stimulation (‘Novel 1’ vs. ‘Novel 2’, based on data in G, I, K). Unpaired two-tailed *t*-tests; Control vs. VIP-ArchT (p = 0.68), Control vs. VIP-ChrimsonR (p = 0.22) and VIP-ArchT vs. VIP-ChrimsonR (p = 0.16). **N)** Population vector correlations using optogenetically stimulated track segments (‘Novel 2’ vs. ‘Novel 3’, based on data in G, I, K). Unpaired two-tailed *t*-tests; Control vs. VIP-ArchT (p = 0.89), Control vs. VIP-ChrimsonR (p = 0.54) and VIP-ArchT vs. VIP-ChrimsonR (p = 0.41).

Mice were first familiarized with a virtual environment for 1-2 weeks before being introduced to a novel environment. To dissect the impact of VIP modulation on different stages of place field formation, we designed a trial structure with four distinct blocks (**Figure 2C**):

1. **Familiar environment (30 trials):** Baseline activity in a well-learned environment.
2. **Novel environment *with* VIP manipulation (‘Novel 1’, 20 trials):** Assess the impact of VIP manipulation on initial place field formation.
3. **Novel environment *without* manipulation (‘Novel 2’, 20 trials):** Allow natural place field development in the novel environment.
4. **Novel environment *with* VIP manipulation reintroduced (‘Novel 3’, 20 trials):** Examine the effect of VIP manipulation on more established place fields.

This design allowed us to assess the role of VIP neurons in both the initial induction of new place fields and their influence on already established place fields.

In control mice, transitioning to the novel environment induced robust place cell remapping, evident in the altered place fields of individual neurons and a change in the overall population activity (**Figure 2D, E**, 26.5 ± 4.3 % of cells were place cells in the familiar environment, and 27.8 ± 4.8 % in the novel environment, mean ± SEM, 6 mice, 11 sessions). This change reflected the formation of a new spatial representation in response to the novel environment. In control mice, place cells formed rapidly during initial novelty exposure (‘Novel 1’) and remained stable in subsequent trial blocks (‘Novel 1,2 & 3’; **Figure 2F**). Population vector correlations across trial blocks (‘Novel 1’ vs. ‘Novel 2’ and ‘Novel 2’ vs. ‘Novel 3’) remained consistent, indicating stable place fields (**Figure 2G**, summarized in **Figure L-N**).

However, inhibiting VIP neuronal activity with ArchT during the initial exposure to the novel environment (**‘**Novel 1’) markedly suppressed the emergence of new place cells (**Figure 2H**) and disrupted the stability of the population code, as evidenced by reduced population vector correlations (**Figure 2I**, 5 mice, 5 sessions). This suppression was spatially localized to the optogenetically stimulated track region (**Figure 2I, L, M**). Importantly, VIP inhibition did not affect the stability of established place fields (‘Novel 3’, **Figure 2I, N**), indicating a specific role for VIP neurons in gating the early stages of place field formation. Conversely, enhancing VIP activity with ChrimsonR slightly but significantly facilitated rapid place cell formation during the initial novelty exposure (‘Novel 1’), but had no effect on established place fields (‘Novel 3’) (**Figure 2J, K, L-N**, 4 mice, 4 sessions). These results demonstrate that VIP interneurons regulate the induction of new place representations in response to novelty but have little affect on established place fields.

### VIP neurons promote place field induction and stabilization, and affect behavioral timescale plasticity

We next examined how VIP modulation influences place cell dynamics, focusing on place fields in the optogenetically stimulated track region. Inhibiting VIP activity during initial novelty exposure (‘Novel 1’) delayed place field formation, but this delay was rapidly compensated when inhibition was removed, allowing normal development (**‘**Novel 2’, **Figure 3A**). The delay in place cell formation was reflected in the time constant of place field formation, which increased from τ = 5.9 trials in control mice to τ = 24.5 trials in VIP-ArchT mice (**Figure 3B**). In contrast, activating VIP cells with ChrimsonR had less effect on the speed of place cell formation (τ = 2.8 trials, **Figure 3A, B**).

**Figure 3.**
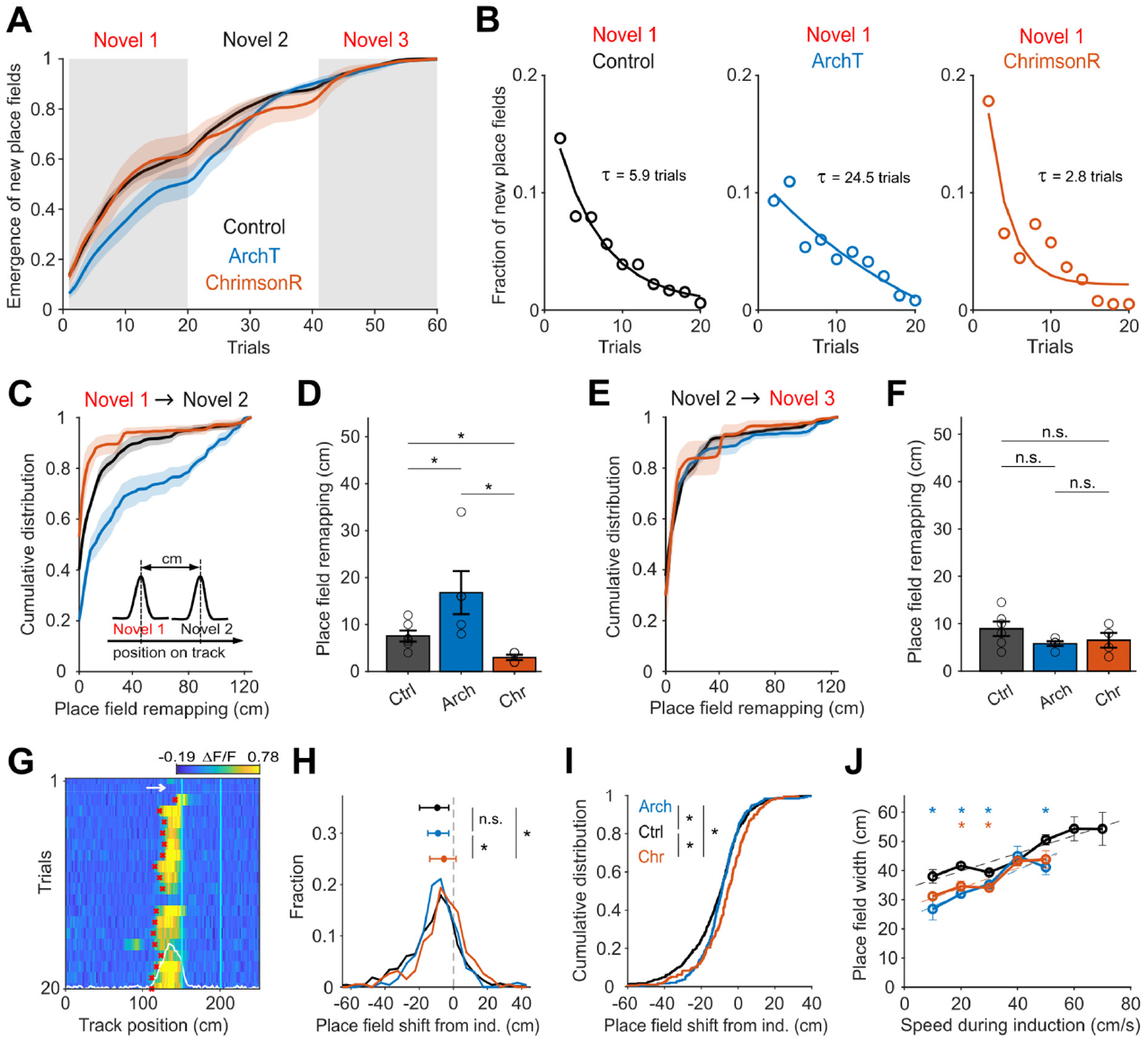
VIP neurons promote place field induction and stabilization, and affect behavioral timescale plasticity. **A)** Fraction of newly formed place fields in the novel environment across trial progression (mean ± SEM, Control: 6 mice, 11 sessions; VIP-ArchT: 5 mice, 5 sessions; VIP-ChrimsonR: 4 mice, 4 sessions). Two-sample Kolmogorov-Smirnov tests: VIP-ArchT vs. Control (Novel 1: p = 0.0082; Novel 2: p = 0.28; Novel 3: p = 0.28) and VIP-ChrimsonR vs. Control (Novel 1: p = 0.97; Novel 2: p = 0.0082; Novel 3: p = 0.5). **B)** Dynamics of place field formation during the Novel 1 trial block. Based on the data in (A); Each data point is the average fraction per 2 trial bins across all sessions. Exponential decay fits: Control (τ = 5.91 trials), VIP-ArchT (τ = 24.51 trials), VIP-ChrimsonR (τ = 2.76 trials). **C)** Cumulative distribution curves (mean ± SEM) of position tuning remapping between Novel 1 and Novel 2 trial blocks. VIP-ArchT vs. Control: Kolmogorov-Smirnov p = 2.6 x 10^-8^; Wilcoxon rank-sum p = 6.3 x 10^-6^, VIP-ChrimsonR vs. Control: Kolmogorov-Smirnov p = 0.0049; Wilcoxon rank-sum p = 0.0012. Control: 6 mice, 11 sessions; VIP-ArchT: 5 mice, 5 sessions; VIP-ChrimsonR: 4 mice, 4 sessions. **D)** Median place field remapping between Novel 1 and Novel 2 environments. VIP-ArchT mice showed increased remapping (p = 0.031, 5 mice), while VIP-ChrimsonR mice showed decreased remapping (p = 0.0092, 4 mice) compared to Control (6 mice). Remapping was also significantly different between VIP-ArchT and VIP-ChrimsonR groups (p = 0.017, one-sided t-test). **E)** Cumulative distribution curves (mean ± SEM) of position tuning remapping between Novel 2 and Novel 3 trial blocks. VIP-ArchT vs. Control: Kolmogorov-Smirnov p = 0.0013; Wilcoxon rank-sum p = 0.33, VIP-ChrimsonR vs. Control: Kolmogorov-Smirnov p = 0.009; Wilcoxon rank-sum p = 0.097. Control: 6 mice, 11 sessions; VIP-ArchT: 5 mice, 5 sessions; VIP-ChrimsonR: 4 mice, 4 sessions. **F)** Median place field remapping between Novel 2 and Novel 3 environments was not significantly different in VIP-ArchT mice (p = 0.95, 5 mice) and in VIP-ChrimsonR mice (p = 0.16, 4 mice) compared to Control (6 mice). Remapping was also not significantly different between VIP-ArchT and VIP-ChrimsonR groups (p = 0.68, one-sided two-sample t-test). **G)** Example of a newly formed place field. Arrow indicates the induction lap. Red asterisks show the onset position in each trial when the place field was active. Note the shift of the onset position compared to the induction trial. White trace shows the mean activity of the place cell. Vertical lines indicate reward locations. **H)** Distribution of place field onset shifts relative to the induction lap in the Novel 1 trial block. Top: Median ± 25% and 75% quartiles for Control (745 place fields), VIP-ArchT (242 place fields), and VIP-ChrimsonR (191 place fields). Comparisons: VIP-ArchT vs. Control (p = 0.11), VIP-ChrimsonR vs. Control (p = 7.4 x 10^-6^), and VIP-ArchT vs. VIP-ChrimsonR (p = 0.0028; two-sided Wilcoxon rank-sum test). **I)** Cumulative distribution of place field onset shifts (same data as (H)). Control vs. VIP-ArchT (p = 0.02); Control vs. VIP-ChrimsonR (p = 0.00027); VIP-ArchT vs. VIP-ChrimsonR (p = 0.0022; two-sample Kolmogorov-Smirnov test). **J)** Place field width as a function of running speed during the induction trial. Data are binned into 10 cm/s bins. The number of place fields in each bin for each group were: Control, n = 106 ± 40; VIP-ArchT, n = 48 ± 20; VIP-ChrimsonR, n = 38 ± 11, mean ± SEM across bins, bins with fewer than 2 data points were excluded. Dashed lines show linear fits for Control (r^2^ = 0.89, p = 0.0014), VIP-ArchT (r^2^ = 0.83, p = 0.031), ChrimsonR (r^2^ = 0.87, p = 0.022). Asterisks indicate a statistically significant difference between VIP-ArchT or VIP-ChrimsonR and Control mice (Wilcoxon rank-sum test, two-sided). P-values for bins 10-50: VIP-ArchT vs. Control: p = 0.013, p = 6.1 x 10^-8^, p = 0.017, p = 0.644, p = 0.0087; and for VIP-ChrimsonR vs. Control: p = 0.20, p = 0.0064, p = 0.0048, p = 0.93, p = 0.062.

Silencing VIP cells during initial exploration (‘Novel 1’) increased place field shifts upon inhibition removal (‘Novel 2’; **Figure 3C, D**), suggesting that VIP activity stabilizes place fields and reduces remapping. Enhancing VIP activity with ChrimsonR reduced place field shifts, indicating more stable place fields (**Figure 3C, D**). These effects were specific to the initial encoding phase, as VIP modulation had no effect on place field shifts in later trial blocks (‘Novel 2 & 3’; **Figure 3E, F**). These results suggest that VIP interneurons accelerate place field formation and enhance stability during early spatial learning.

A subset of hippocampal place cells are formed according to a recently discovered plasticity rule called BTSP ^11,12,40,41^. To investigate how VIP cells influence BTSP, we examined the two key features that characterize it: First, place fields exhibit a forward shift on subsequent trials following the place field induction trial (**Figure 3G**). Second, both the extent of this shift, as well as the place field width, are determined by the animal’s running speed during the induction trial.^11–13^. We found that both silencing and enhancing VIP cells reduced the forward shift of place fields (**Figure 3H, I**) and led to an overall smaller place field width (**Figure 3J**). These results suggest that VIP activity during place field induction regulates key features of BTSP.

### Balanced VIP neuronal activity enhances learning in novel environments but does not affect performance in familiar ones

Having established that VIP cells regulate place field plasticity in novel environments, we next investigated their role in spatial learning. In familiar environments, expert mice exhibited clear signs of learning, displaying anticipatory slowing and spatially constrained licking behavior near the reward locations (**Figure 4A, B**). However, when introduced to a novel environment with new reward locations, this learned behavior disappeared; mice did not adjust their speed, and their licking was no longer localized to the reward zones (**Figure 4E, F**, 5 Control mice, 7 sessions).

**Figure 4.**
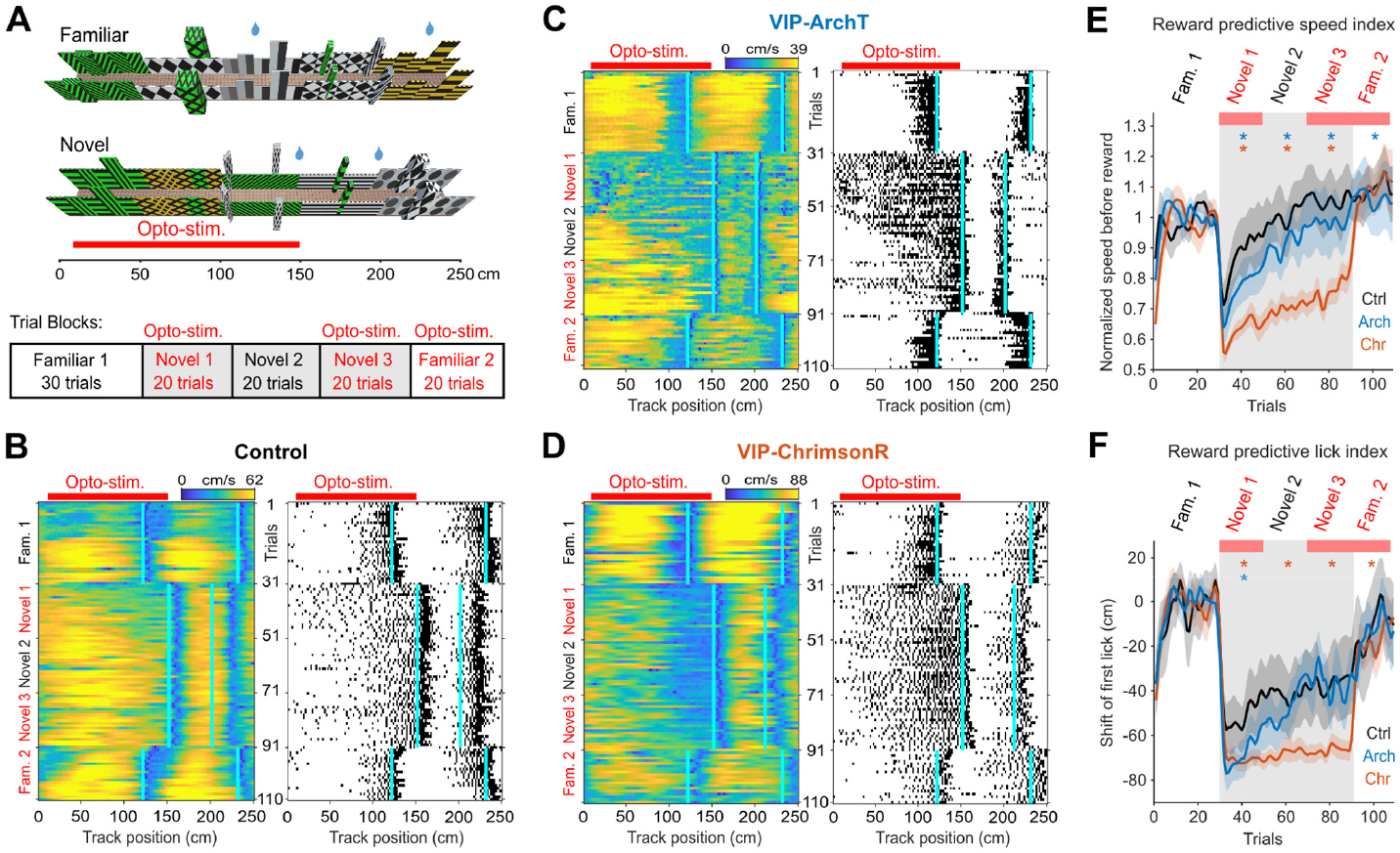
Balanced VIP neuronal activity enhances learning in novel environments but does not affect performance in familiar ones. **A)** Example familiar and novel VR-environment, and the trial block organization. **B-D)** Representative running speed (left) and lick rate (right) traces from a single session for a Control mouse (B), a VIP-ArchT mouse (C), and a VIP-ChrimsonR mouse (D). **E)** Reward predictive speed index. Mean running speed during optogenetic stimulation in the 120 cm before the reward location, normalized to the speed in the familiar block (mean ± SEM, Control: 5 mice, 7 sessions; VIP-ArchT: 7 mice, 10 sessions; VIP-ChrimsonR: 6 mice, 11 sessions). Asterisks indicate significant difference (p < 0.05) between Control and opsin-expressing mice (two-sided Wilcoxon rank-sum test). P-values for novel 1-3 and familiar 2, respectively: VIP-ArchT vs. Control: p = 6.5 x 10^-6^, p = 1.3 x 10^-6^, p = 9.8 x 10^-5^, p = 0.0055; and for VIP-ChrimsonR vs. Control: p = 4.0 x 10^-29^, p = 8.2 x 10^-41^, p = 5.7 x 10^-36^, p = 0.46. **F)** Reward predictive lick index. Change of the first lick position relative to the familiar environment, analyzed within the optogenetically stimulated part of the corridor (120 cm range before the reward location), mean ± SEM, black: Control: 5 mice, 7 sessions; VIP-ArchT: 7 mice, 10 sessions; VIP-ChrimsonR: 6 mice, 11 sessions). Asterisks indicate a significant difference (p < 0.05) between Control and opsin-expressing mice (two-sided Wilcoxon rank-sum test). P-values for novel 1-3 and familiar 2, respectively: VIP-ArchT vs. control: p = 1.3 x 10^-8^, p = 0.3, p = 0.92, p = 0.06; and for VIP-ChrimsonR vs. Control: p = 2.4 x 10^-10^, p = 1.4 x 10^-12^, p = 4.4 x 10^-11^, p = 0.0064.

Silencing VIP activity during this learning phase impaired their ability to learn the new reward locations, and, remarkably, enhancing VIP activity led to even more pronounced learning deficits (**Figure 4C, D, E, F**, 7 VIP-ArchT mice, 10 sessions; 6 VIP-ChrimsonR mice, 11 sessions). In contrast, manipulating VIP activity in the familiar environment had much less effect on performance (**Figure 4C, D, E, F**). These findings indicate that a balanced level of VIP activity is essential for efficient spatial learning in novel environments, while established spatial memories are more resistant to VIP modulation.

## Discussion

We found that exposure to a novel environment triggered a transient increase in CA1 VIP neuron activity, mirroring arousal changes measured by pupil diameter (**Figure 1F, G**). Individual VIP neurons exhibited diverse response dynamics, with some activating immediately upon novelty exposure and others responding more gradually (**Figure 1E**). The population response suggests that environmental novelty triggers arousal and learning, mediated in part by CA1 VIP interneurons.

VIP-mediated inhibition primarily targets SOM-expressing interneurons, particularly OLM cells in hippocampal area CA1 ^33,35,65^. OLM cells exert powerful inhibitory control over the distal tuft dendrites of pyramidal cells ^37,38^, the same dendrites that receives excitatory input from layer 3 of the entorhinal cortex (EC3), which is thought to provide an instructive signal for new place field formation ^13,66^. Because coincident activation of EC3 and CA3 inputs is required for generating BTSP, a plasticity mechanism important for place field formation ^13,67,68^, OLM cells are strategically positioned to gate the induction of new place fields. Our findings reveal that VIP-mediated disinhibition directly modulates BTSP properties, with both silencing and enhancing VIP interneuron activity reducing the forward shift of place fields and decreasing place field width (**Figure 3H-J**). These results indicate that VIP interneuron activity during place field induction controls key features of BTSP. This transient disinhibition during novelty will reduce OLM-mediated suppression of dendritic excitability, facilitating the integration of coincident EC3 and CA3 inputs and increasing the likelihood of reaching the threshold for dendritic spike generation and plasticity induction. Consistent with this model, a recent key paper shows that OLM cells are transiently silenced in novel environments, and their artificial activation reduces remapping ^39^.

We propose that novelty-induced arousal triggers the release of neuromodulators, including acetylcholine, noradrenaline, dopamine, and serotonin, some of which are known to affect spatial map formation ^69–72^ and strongly excite VIP interneurons directly ^51,57,60,73^. This will lead to transient disinhibition of pyramidal cells, thereby facilitating the formation of new place maps. This is consistent with recent work showing that VIP neurons respond to rewards and enhance the representation of reward locations in the hippocampus ^43^. However, our findings indicate that VIP-mediated disinhibition alone is insufficient for remapping. Optogenetic activation of VIP neurons in a familiarized novel environment had minimal effects on remapping, place field stability, and behavior (**Figures 2N, 3E-F**, and **Figure 4**). This suggests that neuromodulators are also required and act not only on VIP neurons but also directly on pyramidal cells and other interneuron populations ^74,75^. The coordinated actions of these neuromodulators likely enhance dendritic excitability and facilitate plasticity mechanisms such as BTSP, underscoring the importance of combined VIP-mediated disinhibition and broader neuromodulatory influences for effective remapping.

In the hippocampus, VIP interneurons can be classified into four subtypes: three disinhibitory types (VIP+/Calretinin-, VIP+/Calretinin+, VIP+/muscarinic R2+) and one basket-type cell (VIP+/CCK+) that directly inhibits pyramidal cells ^18,76^. VIP+/CCK+ basket cells ^26,43,77^ and VIP+/muscarinic R2+ cells ^26,35^ are suppressed during locomotion but active during rest, while disinhibitory VIP subtypes are active during locomotion and primarily target OLM cells. Consequently, inhibiting VIP cells during locomotion predominantly affects the disinhibitory subtypes, as the VIP+/CCK+ basket cells are already inactive. In contrast, ChrimsonR activation would engage all VIP subtypes, potentially explaining the more pronounced effects of activation on learning processes (**Figure 4E, F**).

The role of VIP-mediated disinhibition in shaping sequential activation patterns extends beyond the hippocampus. In the primary motor cortex, neural sequences encoding motor actions are dynamically modulated during learning ^78^. Similar to our findings in the hippocampus, inhibiting VIP cells in the motor cortex prevents the adaptation of these neuronal sequences and impairs the acquisition of new motor skills ^78^. Similarly, activating SOM cells during well-learned motor tasks has no effect ^78,79^, indicating that disinhibition is selectively engaged during learning. Further paralleling our work, both optogenetic up- and downregulation of SOM cell activity impair motor skill learning ^79^ (**Figure 4**), suggesting that a precise balance of dendritic inhibition is crucial. This balance has been proposed to be critical for activity-dependent dendritic spine reorganization ^79^, likely mediated by CaMKII-dependent synaptic plasticity ^78,80^. Optogenetic activation or suppression of SOM cells impaired learning new motor skills by destabilizing and hyper-stabilizing spines, respectively ^79^. Additionally, artificial VIP activation may induce widespread, nonspecific dendritic spikes, potentially causing memory interference ^81,82^. Together, these striking parallels between hippocampal and motor cortical circuits—despite their distinct roles in spatial cognition and motor control— highlight that VIP-mediated regulation of plasticity is a fundamental and widespread mechanism for neural circuit adaptation.

## Acknowledgments

This work was funded by the European Research Council (ERC starting grant #639272), the Research Council of Norway #231495, #276047, and #274306), and a MSCA fellowship to Nora Lenkey (#789962). We thank Kristin Larsen Sand, Eivind Hennestad and the IMB mechanical and electrical workshop for their technical assistance.

Finally, we thank the members of the Vervaeke lab for providing critical comments that significantly improved a draft of the manuscript.

## Author contributions

Conceptualization, M.N., N.L. and K.V.; methodology, M.N., N.L. and K.V; investigation, M.N., N.L.; software, M.N.; formal analysis, M.N.; supervision, K.V.; writing – original draft, M.N., N.L. and K.V.; writing – review & editing, M.N., N.L. and K.V; funding acquisition, N.L. and K.V.

## Methods

### Resource availability

Lead contact: Further information and resource requests should be directed to and will be fulfilled by the lead contact Koen Vervaeke.

### Materials availability

This study did not generate new unique reagents.

### Data and code availability

- The lead contact will share the mouse behavior and Ca2+ imaging data reported in this paper upon request.
- All original code will be deposited on GitHub and is publicly available as of the date of publication.
- Any additional information required to reanalyze the data reported in this paper is available from the lead contact upon request.

### Experimental model and subject details

We used either VIP-IRES-Cre mice ^58^ (JAX #010908, 6 mice for imaging VIP cells expressing GCaMP6s) or Thy1-GCaMP6s mice ^83^ (GP 4.3 line #024275 from JAX) crossed with VIP-IRES-Cre mice (7 mice for VIP-ArchT x Thy1-GCaMP, 6 mice for VIP-ChrimsonR x Thy1-GCaMP6s, 6 mice for no-opsin (Thy1-GCaMP6s) control experiments, including both imaging and behavior experiments). The VIP-Cre mice expressing GCaMP6s, ArchT and ChrimsonR were also used for a previous study ^64^. Mice were 4.09 ± 0.38 months old at the time of surgery (mean ± SEM, 25 mice were used in total), and we used both male and female mice. We housed mice in groups with 1-4 littermates on a reversed 12-hour light / 12-hour dark cycle, and we carried out experiments during their dark phase. Mice were kept in a humidity and temperature-controlled environment. For training and experiments, they were kept on a water restriction regime as described previously ^84^ and received water drop rewards throughout the task. All procedures were approved by the Norwegian Food Safety Authority (FOTS 6590, 7480, 19129, 30014), and experiments were performed in accordance with the Norwegian Animal Welfare Act.

## Methods details

### Virtual reality behavior setup, software for behavior control and acquisition

We used a custom-built virtual reality behavior setup. Mice run on a Styrofoam running wheel with a circumference of 157 cm (50 cm diameter, 10 cm wide; Bakedeco, Brooklyn, New York), which we rubber-coated to achieve a highly durable and washable surface. First, we filed the surface with the help of a lathe, then a layer of plastic wrapping tape was applied on the wheel to protect the Styrofoam from the solvent, finally a thin layer of rubber spray (Plasti Dip) was spread on the surface in 2-3 steps. A rotary encoder (Omron E6B2-CWZ1X) attached to the wheel registered the locomotion of the mouse, this signal controlled the movement through the VR-corridors, projected on 2 monitors that were placed in a V-shape in front of the mouse. In total, we used 4 different VR-environments (one familiar and 3 novel environments, implemented in Unity). Each VR-corridor was 250 cm long, and circular (the teleportation from the end to the starting point was seamless), and each hosted 2 separate reward locations. The behavior was automated using custom-written LabVIEW software (National Instruments, 2017) to communicate with Unity, control all actuators, sensors, and data acquisition, record behavioral parameters (3 kHz), and trigger the microscope to start and stop recording. The lick-port signals, rotary encoder signals, and two-photon frame clock were acquired using a DAQ (X-Series, PCIe 6321, National Instruments). We used the two-photon frame clock signal to synchronize the two-photon images with the recorded signals via the DAQ. The behavior setup was positioned under a custom-built two-photon microscope in a light-tight enclosure built from Thorlabs parts (25 mm optical rails, Thorlabs, XE25L48), blackout hardboard (Thorlabs, TB4) and blackout fabric (Thorlabs, BK5). We controlled water reward delivery with a solenoid valve and a custom-made lick port ^84^ to detect licks.

### Pupil tracking

For tracking the pupil diameter an infrared-sensitive camera (Basler dart 1600-60um) was mounted to the head-bar holder frame of the behavior setup, below the horizontal level of the left eye to avoid blocking the vision of the monitors. The frame rate was 31 or 60 fps, and the frames were acquired using custom-written LabVIEW code (National Instruments) along with the frame-clock that allowed for subsequent alignment with the behavior and 2P data. The pupil was back-illuminated by the infrared light from the two-photon laser during imaging experiments, which created a high contrast image of the pupil. During post-processing, we measured the pupil position and diameter using custom written MATLAB scripts.

### Virus injections

We used AAV1-syn-flex-GCaMP6s (Addgene #100845) for imaging VIP cells, and AAV5-Syn-flex-rc[ChrimsonR-tdTomato] (Addgene, #62723) or AAV5-CAG-flex-ArchT-tdTomato (Addgene, #28305) for optogenetic modulation of VIP cells. In some cases, the virus was diluted with sterile PBS to get a virus titer of 10^12^. Virus injections in area CA1 were done at least one week before the window implantation. We injected the virus to one location in the dorsal area CA1: AP - 2.0, ML + 1.2, at 1.2 and 1.05 mm from the brain surface. For optogenetic activation of VIP cells, we injected 2 x 100 nl of AAV5-Syn-flex-rc[ChrimsonR-tdTomato], and for optogenetic inactivation of VIP cells, we injected 2 x 100 nl of AAV5-CAG-flex-ArchT-tdTomato. For Calcium imaging of VIP cells, we injected 100 nl per injection site of AAV1-syn-flex-GCaMP6s. Animals were used for experiments between 1-4 months after virus injection. The GCaMP6s, as well as the opsin expression, were always checked with a 2P microscope before we started experiments.

### Surgery procedures

Surgery was carried out under isoflurane anesthesia (3% induction, 1-1.5% maintenance) while maintaining body temperature at 37 degrees Celsius with a heating pad (Harvard Apparatus). We delivered a subcutaneous injection of 0.1 mL Marcaine (bupivacaine 0.25% m/V in sterile water) at the scalp incision site. Virus injections into CA1 were performed at least one week before the craniotomy. After removing the scalp, we implanted a titanium head bar and made a circular cranial window with a 2.5 mm diameter using a dental drill. The center of the window was AP -2.0 mm, ML 1.5 mm from bregma. After drilling a 2.5 mm craniotomy, we removed cortical tissue and most of the corpus callosum above dorsal CA1 using a suction device (∼40 bar). We implanted a metal ring (diameter 2.5 mm, height 1.5 mm) glued to a thin glass window at the bottom (#1.5 coverslip glass) with optical adhesive (Norland Optical Adhesive; Thorlabs NOA61). We affixed the metal ring to the skull with cyanoacrylate glue, and we covered the exposed skull with dental cement outside the metal ring ^85^. We administered postoperative analgesia (Temgesic, 0.1 mg/kg) subcutaneously, and animals were monitored for 2-3 days after surgery for any sign of distress or pain, and additional analgesia was administered if needed.

### Water restriction and animal training

Mice were water-restricted after a 10-day post-operative recovery period ^84^. Animals received 1-1.5 ml water per day while we monitored their body weight to ensure they maintained at least 80% of their initial body weight. During the first week of water restriction, we handled mice daily in dim light conditions to gradually habituate them to the experimenter and to the head fixation. During the training process, mice first learned to run head-fixed without the VR on a blank polystyrene wheel, receiving randomly delivered water rewards (2-5 µl/drop). We decreased the reward frequency and increased the session length gradually until the mice reached the performance required for the VR task (∼100 trials in ∼20 minutes with 125cm reward frequency). This initial training phase took typically 5-7 training sessions, then we transitioned mice to the rubber coated experimental wheel and introduced the VR-task. We trained mice for minimum 5 days in the familiar environment, under the microscope before the experiments.

### Two-photon Calcium imaging and optogenetic manipulations

We used a custom-built two-photon microscope (INSS) designed to provide enough space under the objective to accommodate the large running wheel and monitors for the virtual reality system. We acquired images at 31 Hz (512 x 512 pixels) using the open-source acquisition software SciScan (LabVIEW, National Instruments). The excitation wavelength was 930 nm using a MaiTai DeepSee ePH DS laser (SpectraPhysics). The average power measured under the objective (N16XLWD-PF, Nikon) was 30–90 mW. Photons were detected using GaAsP photomultiplier tubes (PMT2101/M, Thorlabs). The primary dichroic mirror was a 700 nm LP (Chroma), and the photon detection path consisted of a 680 nm SP filter (Chroma), a 565 nm LP dichroic mirror (Chroma), and to prevent the intense red light from entering the photomultiplier of the green channel, we mounted two optical density 6 filters (#ET510/80, Chroma) in series after the secondary dichroic, slightly angled in relation to each other to achieve a higher optical density attenuation (#T565lpxr, Chroma). The typical FOV size was 628-742 x 628-742 µm for VIP cell imaging and between 583-680 x 583-680 µm for pyramidal cell imaging. We recorded VIP cells or pyramidal cells at 80–150 µm depth from the top of CA1 (alveus).

In the optogenetic trial blocks (in experiments related to Figures 2-4), the optogenetic stimulation was turned on between positions ∼10 and 150 cm in each trial. To let some recovery time for the circuit, and as an internal control, the remaining part of the track (positions 150-250 cm, that included the consumption of both water reward in the new environments), remained unstimulated. The light stimulation lasted typically for 5-8 seconds, with a cut-off time of 10 seconds. For the optogenetic track segments, we included data between 20-150 cm to account for variability of stimulus onset due to the behavioral software running on a soft real time operating system. For ArchT stimulation we used a square shaped light pulse, whereas for ChrimsonR stimulation we used a 20 Hz sinusoid light pulses. To implement simultaneous two-photon Ca^2+^ imaging and optogenetic modulation, we mounted a red laser diode (Oclaro, 638 nm 700 mW) in the position where normally the red channel photomultiplier would be located. To make the beam shape more circular, we focused the light into an optical fiber (Thorlabs, #M15L01) using a fiber coupler with a collimation lens (Thorlabs, #PAF2S-11B) ^86^. The beam was then focused on the objective back aperture such that it diverged when entering the brain. The diameter of the beam was approximately 2.6-3 mm, the intensity was 6.4-7.2 mW/mm^2^. Since the red light used for optogenetics could spread through the brain and stimulate the back of the retina, we used a similar-wavelength masking light positioned in front of the eyes (Thorlabs, LED630E and LED630L). This masking light (20 Hz) was on during the entire session.

### Image processing

Images were registered using a combination of custom and published code. Images were first de-stretched to correct for distortions resulting from the sinusoidal speed profile of the resonance scan mirror of the microscope. Next, rigid and non-rigid motion correction was performed using custom-written scripts using Flow-Registration ^87^. For image segmentation, somatic regions of interest (ROIs) were detected using a custom auto-segmentation method followed by manual curation after visual inspection of each ROI. To correct neuropil contributions, a doughnut-shaped ROI of the surrounding neuropil was automatically created for each ROI by dilating the neuropil ROI four times larger than the original ROI. If the soma ROI or the neuropil ROI overlapped with another soma ROI, that area was excluded from the ROI. Then, pixel values were averaged for each ROI per imaging frame. ROI fluorescence changes were defined as a fractional change DF/F according to DF/F(t) = (F(t)-F0)/F0, where F0 is the baseline activity defined as the 20th percentile of the ROI fluorescence signal. We calculated DF/F(t) values for all ROIs and related neuropil areas, which were subtracted from the ROI DF/F(t) values, and a correction factor was added to ensure that the soma DF/F remained positive. DF/F(t) signal was used for further analysis. Importantly, F0 was a single value applied to the entire session such that it would not correct for baseline fluorescence changes during optogenetic manipulations. ROIs with signal to noise ratio <2 were excluded from further analysis.

### Quantification and statistical analysis

We analyzed DF/F and behavioral time series with MATLAB. The behavioral time series were first binned to match the two-photon microscope frame rate (31 Hz) by finding the behavioral data sample closest in time to the imaging frame or, for some signals, taking the average of some samples around that sample.

### Classification of VIP-GCaMP neuron activity in novel environment

We took 10 trials before and after the transition between environments and calculated the mean activity of each trial. To test whether the cell’s activity is upregulated in the novel environment we used a one-sided Wilcoxon rank-sum test.

### Thy1-GCaMP recordings

For each session imaging Thy1-GCaMP6s neurons, we assessed two quality control criteria: (1) mice had to maintain task engagement shown by consuming water rewards throughout all trial blocks, (2) mice had to have a stable neuronal representation of the familiar environment, that has been assessed by population vector cross correlation of all the neurons’ activity between odd and even trials (see below for more details). Sessions with < 0.8 correlation coefficient were excluded from the analysis. Based on these criteria, we included n = 20 sessions (control: n = 11, VIP-ArchT: n = 5, and VIP-ChrimsonR: n = 4) in the analysis.

### Place cell classification

To identify position tuned cells in each trial block, we used a bootstrapping approach. First, we binned the DF/F signals of each ROIs according to the position in the VR environments by averaging the activity into 2 cm bins. Next, we calculated a position tuning curve by averaging the activity of all trials within the trial block. We compared the peak amplitude to a reference distribution calculated the same way, after random shuffling the same dataset for 1000 iterations. The shuffling was done by circular shifting the activity within each trial. To test the reliability, we split the trials into odd and even groups and calculated the tuning curves for each, then calculated the Pearson correlation coefficients. We compared this data with the 1000 times shuffled distribution of the original data. We considered a cell position tuned if it passed the >95% threshold for both parameters. In addition, to be classified as a place cell within a trial block, neurons had to have at least one, but not more than three place fields.

### Place field detection

The place field detection algorithm was optimized to find newly generated place fields. First, we generated a separate position tuning curve for each trial block by averaging the trials (except for the first block where we used the last 20 trials to match the trial count of the other blocks). We used the tuning curves to identify potential place fields using the ‘findpeaks’ function (MATLAB, Signal Processing Toolbox) with a minimum peak height (defined as the input ‘prominence’ parameter) of 5 % of the highest activity bin of a ROI. Next, we looked for significant activity peaks (prominence is > 2.5 x SD of the activity bins of the entire session) on trial-by-trial basis within the width of the potential place field (defined as peak ± half width) and analyzed these significant activity peaks within a moving window of 5 trials. The *induction lap* was defined as the first trial with a significant peak that is followed by two more active trials within the next 4 trials. We considered a place field active until the activity dropped below the 60% reliability threshold within the moving window, where we determined the last active trial.

A potential field had to fulfill the following criteria to be accepted as a place field: 1) place field width is between 6 - 80 cm; 2) 60% reliability as described above; 3) maintain its activity for minimum 5 trials between the induction and the last active trial. Note that the moving window was not restricted to the trials within the tested block, i.e. when a potential induction lap was detected in the last trial of a block, the moving window was shifted until the 4^th^ trial of the subsequent trial block so that this field could still meet the criteria.

For the BTSP analysis we also defined the *place field onset shift. W*e detected the onset positions of the significant activity peaks in each trial, as the first position bin before the peak where the activity crosses the threshold of 2 x SD of the binned ROI activity, and calculated the onset shift relative to the induction trial as the mean of all following trials. For the *induction speed*, we determined the running speed as the mean speed at the induction peak bin and the preceding 4 position bins.

### Population vector correlation

To correlate the population activity of two trial blocks, for each we generated matrices from the position tuning curves including all place cells of the novel trial blocks. We calculated the position tuning curves as the mean binned DF/F signals of all trials within a trial block. We used Pearson correlation on these matrices to gain a matrix of correlation coefficients of each pair of position bins. To quantify the data, we calculated the maximum correlation coefficient values for each position and took the median value of either the optogenetically stimulated (20 - 150 cm) or the non-stimulated (150 – 250 cm) segment of the track.

### Peak position remapping

To quantify remapping of place fields between two trial blocks (Novel 1 vs. Novel 2, and Novel 2 vs Novel 3, Figure 3), for cells that we classified as place cell in both trial blocks, we calculated the absolute minimum distance between the peak position of the place fields in the trial blocks. When a neuron had multiple place fields, we paired them so that the pair with the minimum distance was selected first, followed by the second smallest and so on. Unpaired fields such as newly formed fields were ignored in this calculation.

### Behavior Data

#### Lick Rate

For each session, we assessed two quality criteria: (1) mice had to maintain task engagement by consuming water rewards throughout all five trial blocks, and (2) the noise level of the lick signal. To reduce false licks caused by high-frequency noise, we applied a low-pass filter (>15 Hz and <15 ms length) to the recorded binary lick signal. Sessions were excluded from analysis in Figure 4 if false licks constituted more than 15% of the total lick count.

#### Running Speed

Running speed was calculated by differentiating the elapsed distance recorded from the rotary encoder, with the resulting signal smoothed using a 100 ms Gaussian smoothing window.

To analyze the behavioral effects of optogenetic manipulations, we used lick and running speed data from a range of 120 cm before the first reward location (0–120 cm in the familiar and 30–150 cm in novel environments). Within this window, we calculated the following measures that allow for direct comparison of changes between groups relative to the familiar environment:

#### Shift of the First Lick

The position of the first lick before the reward was identified in each trial, then we subtracted the mean of the last 20 trials of the familiar trial block (Figure 4E).

#### Normalized Running Speed

The mean running speed of each trial was normalized by dividing it by the mean running speed of the last 20 trials of the familiar trial block (Figure 4F).

To increase statistical power, we included up to two sessions per mouse in the dataset (control: n = 7 sessions, VIP-ArchT: n = 10 sessions, VIP-ChrimsonR: n = 11 sessions).

### Statistics

If the dataset did not pass the Kolmogorov-Smirnov normality test, non-parametric statistical tests were used (two-sided Mann-Whitney U test for independent samples). Significance was set at p < 0.05. For normally distributed data, and data with sample size n ≤ 10, we used paired or unpaired one- or two-sided t-tests. For comparison of cumulative distributions, we used Two-sample Kolmogorov-Smirnov test. The figures show mean ± SEM unless stated otherwise.

